# Transcriptomic analysis of temporal shifts in berry development between two grapevine cultivars of the Pinot family reveals potential genes controlling ripening time

**DOI:** 10.1101/2021.03.18.436038

**Authors:** Jens Theine, Daniela Holtgräwe, Katja Herzog, Florian Schwander, Anna Kicherer, Ludger Hausmann, Prisca Viehöver, Reinhard Töpfer, Bernd Weisshaar

## Abstract

**Background:** Grapevine cultivars of the Pinot family represent clonally propagated mutants with major phenotypic and physiological differences, such as different colour or shifted ripening time, as well as changes in important viticultural traits. Specifically, the cultivars ‘Pinot Noir’ (PN) and ‘Pinot Noir Precoce’ (PNP, early ripening) flower at the same time, but vary in the beginning of berry ripening (veraison) and, consequently, harvest time. In addition to genotype, seasonal climatic conditions (i.e. high temperatures) also affect ripening times. To reveal possible regulatory genes that affect the timing of veraison onset, we investigated differences in gene expression profiles between PN and PNP throughout berry development with a closely meshed time series and over two separate years.

**Results:** The difference in the duration of berry formation between PN and PNP was quantified to be approximately two weeks under the growth conditions applied, using plant material with a proven PN and PNP clonal relationship. Clusters of co-expressed genes and differentially expressed genes (DEGs) were detected which reflect the shift in the timing of veraison onset. Functional annotation of these DEGs fit to observed phenotypic and physiological changes during berry development. In total, we observed 3,342 DEGs in 2014 and 2,745 DEGs in 2017 between PN and PNP, with 1,923 DEGs across both years. Among these, 388 DEGs were identified as veraison-specific and 12 were considered as berry ripening time regulatory candidates. The expression profiles revealed two candidate genes for ripening time control which we designated *VviRTIC1* and *VviRTIC2* (VIT_210s0071g01145 and VIT_200s0366g00020, respectively). These genes likely contribute the phenotypic differences observed between PN and PNP.

**Conclusions:** Many of the 1,923 DEGs show highly similar expression profiles in both cultivars if the patterns are aligned according to developmental stage. In our work, putative genes differentially expressed between PNP and PN which could control ripening time as well as veraison-specific genes were identified. We point out connections of these genes to molecular events during berry development and discuss potential candidate genes which may control ripening time. Two of these candidates were observed to be differentially expressed in the early berry development phase. Several down-regulated genes during berry ripening are annotated as auxin response factors / ARFs. Conceivably, general changes in auxin signaling may cause the earlier ripening phenotype of PNP.

## Background

*Vitis vinifera subsp. vinifera* (grapevine) belongs to the family *Vitaceae*. With 6,000 to 11,000 cultivars, it is one of the most important perennial crops worldwide [1]. Grapevine fruit development can be divided into two physiological phases, berry formation and berry ripening. Veraison refers to the transition from berry formation to berry ripening, and each of the two phases is represented by a sigmoidal growth curve of development [2]. The progress through development is described by stages referred to as “BBCH stages” (acronym derived from the names of the coordinating institutions involved in stage definition) that have been defined for several crops including grapevine [3, 4]. The first physiological phase is described as berry formation (berry initiation and growth with cell divisions) and lasts from the end of flowering (BBCH71) until ∼60 days later when the majority of berries are touching each other (BBCH79). The developmental stage of veraison (BBCH81) is the end of berry formation and the start of berry ripening [2]. Phenotypically, veraison is the developmental switch when the berries start to soften, accompanied by the onset of accumulation of phenylpropanoids. In red grapevine cultivars, veraison is also indicated by a colour change of the berries that is caused by the beginning of accumulation of anthocyanins, a major class of phenylpropanoids. Members of the well-studied protein superfamily of R2R3-MYB transcription factors (TFs) are considered to be mainly accountable for controlling anthocyanin accumulation [5–7]. Berry ripening starts at veraison and continues until harvest (BBCH89), this phase includes cell enlargement, sugar accumulation and acidity decline. Timing of veraison has also been studied at the level of genetic loci and genomic regions that control this trait. Since it is a quantitative trait influenced by several to many genetic loci, quantitative trait locus (QTL) analyses have been performed. These studies detected a major QTL for timing of the onset of veraison on chromosome 16, combined with a minor QTL on chromosome 18 [8]. By integrating a number of QTL studies, several meta-QTLs connected to genetic control of veraison time were detected, with the most relevant located on chromosome 14, 16, and 18 [9].

Anthropogenic climate change is resulting in successively earlier ripening of grapes with a significant impact on berry quality and consequently the expected flavours of a desired wine style [10]. In addition, the time of veraison and harvest of a given cultivar may differ greatly, driven by regional and/or year-specific differences in weather conditions. Obviously, this calls for a better molecular understanding of the control of ripening time in grapevine. Comparison of different grapevine cultivars grown at the same environmental conditions often uncovers differences in the duration of berry formation, timing of veraison, duration of berry ripening, and ripening time in general. However, the underlying genetic factors are mostly unknown. Previous studies have elucidated how ripening time is affected by internal and external factors. For example, the effect of phytohormones on berry ripening has been widely studied [1]. In general, fruit growth is discussed to be controlled by several phytohormones which play essential roles to trigger or delay ripening processes [11]. Grapevine is a non-climacteric fruit and effects of abscisic acid (ABA) have been investigated in many studies as ABA is considered to trigger ripening [12–14]. Furthermore, it was shown that ABA is involved in controlling leaf senescence [15], responses to drought [16] and pathogen defense [17]. In grapevine, although not as central as in climacteric fruits like tomato (*Solanum lycopersicum*), the phytohormone ethylene is involved in the control of berry ripening [1, 13, 18, 19], while auxin has been shown to induce a delay of ripening [20, 21].

Fruit development of both, dry and fleshy fruits, has been studied very intensively for the obvious reason that fruits are central to human nutrition [22, 23]. The main model system for studies on fleshy fruits is tomato, because of established genetics and molecular biology, access to mutants, and well advanced transgenic approaches to gene function identification [24, 25]. Berry development of grapevines has also been studied intensively [1, 26] and often at the level of the transcriptome. In quite some of the studies, predominantly late berry development stages were sampled to bring the development stage of veraison into the focus [9, 27–29]. In addition, whole berry development was studied with coarse time point distribution [30–34].

To monitor gene expression changes at a high resolution throughout grapevine berry development, starting from flowering until berries are matured, we sampled a comprehensive time series from two Pinot cultivars across two years. The samples were collected from the grapevine cultivar ‘Pinot Noir’ (PN) and the comparably earlier ripening cultivar ‘Pinot Noir Precoce’ (PNP) that is expected to be closely related to PN. The cultivar PNP is listed in the *Vitis* International Variety Catalogue (*V*IVC; [35]) and described to flower at the same time as PN but to reach veraison significantly earlier than PN [36]. Quantitative data for transcript levels, interpreted as values for gene expression, were generated by RNA-Seq. We studied the general course of gene expression patterns throughout berry development in both years and cultivars, and identified a number of differentially expressed genes (DGEs) between PN and PNP prior to veraison. These DEGs can be considered as important candidates for either delaying or pushing forward berry development. Our main aim was the identification of genes controlling the speed of development, to offer an entry point into characterization of the relevant molecular functions in grapevine, and to facilitate future breeding strategies that address traits relevant to, and affected by, climate change.

## Results

### Phenotypical comparison between two Pinot cultivars

To study ripening shifts, we used samples of two closely related grapevine cultivars. The cultivar PNP is an earlier ripening clonal variant of its ancestor PN. Clonal relation of PN and PNP was confirmed by a set of 24 SSR markers that all displayed the identical allele status for both cultivars (Additional file 1: Table S1). To confirm and validate the phenotypic differences between PN and PNP, detailed BBCH developmental stages were determined and documented (Figure 1). PN and PNP display similar phenotypic properties during development and flower (BBCH65) at the same time. However, veraison (BBCH81) is shifted to ∼2 weeks earlier for PNP, and similar shifts were observed in four different years (Table 1). In addition, Figure 1A shows an overview across the time points at which samples were taken. The phenotypic differences between PN and PNP are illustrated in images of developing berries taken between onset of berry formation and veraison (Figure 1B and Additional file 1: Table S2 and Table S3). Veraison (BBCH81) is visible as the onset of anthocyanin accumulation and is detected ∼2 weeks later in PN compared to PNP.

**Figure 1:**
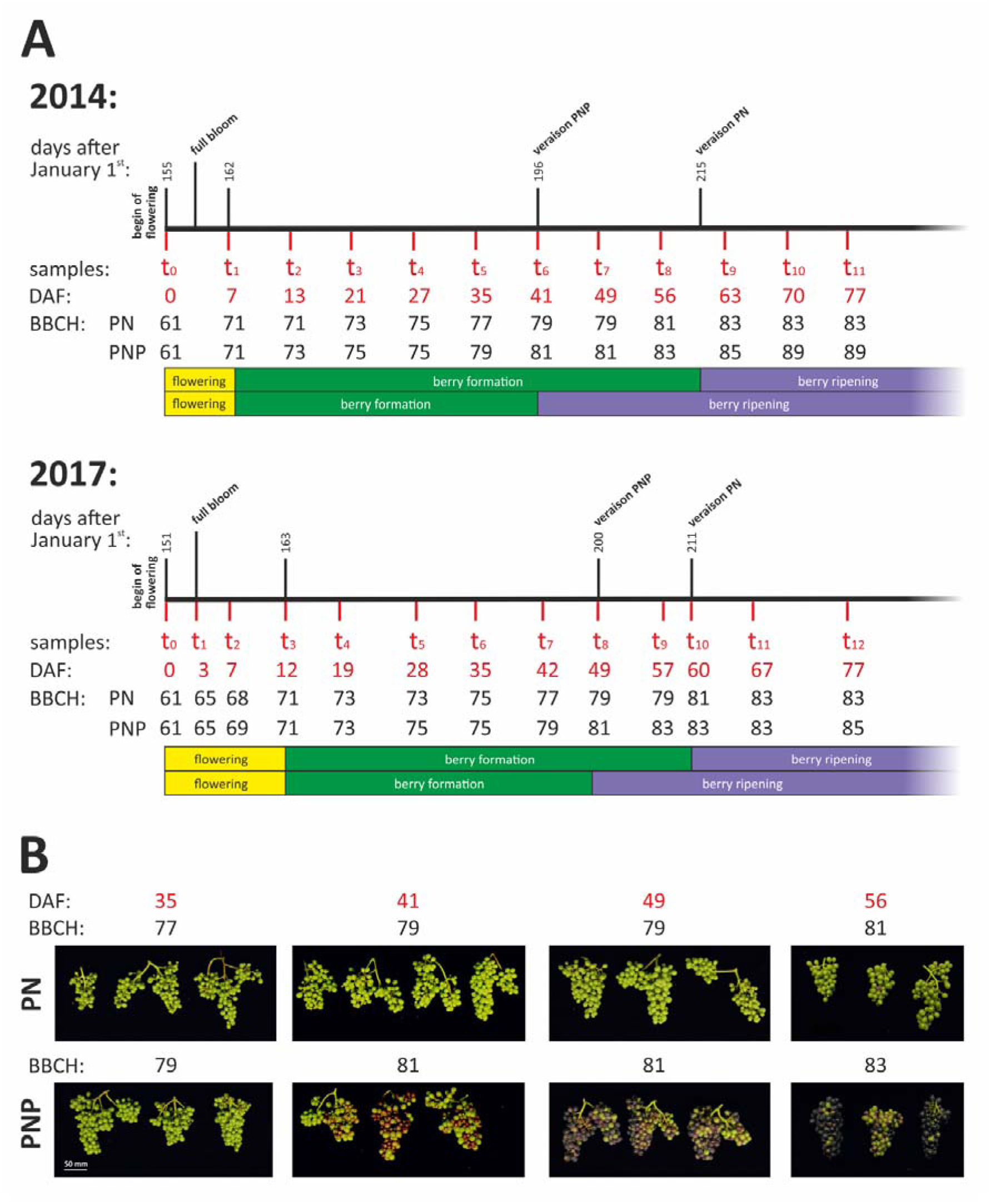
Phenotypical observations and sampling scheme. (A) Sampling time points and days after onset of flowering (DAF) are indicated in red. The developmental stage observed is shown in the BBCH stages [3, 4]. DAF zero (0) is set at BBCH61 (onset of flowering, 10% of flowerhoods fallen [3]). Berry development is depicted schematically and categorized into the phases flowering (yellow), berry formation (green), and berry ripening (purple) for both cultivars. The junction between green and purple indicates veraison (BBCH81). To orient for time of year, numbered days after January 1^st^ are shown. (B) Images of grape bunches and developing berries taken in 2014 are shown to document the differences between PN and PNP. Images were taken 35, 41, 49 and 56 DAF. Scale bar: 50 mm.

**Table 1:** Observed flowering- and berry development shifts between the cultivars PNP and PN in 2014, 2015, 2016 and 2017 at the Geilweilerhof vineyards, Siebeldingen, Germany (in days after January 1^st^).

| Year | Cultivar | Start of flowering (BBCH61) | End of flowering period / Start of berry formation (BBCH71) | End of berry formation / veraison (BBCH81) | Flowering time [Δ days] | Berry formation time [Δ days] |
| --- | --- | --- | --- | --- | --- | --- |
| 2014 | PNP | 155 | 162 | 196 | 0 | 19 |
| 2014 | PN | 155 | 162 | 215 |  |  |
| 2015 | PNP | 159 | 166 | 201 | 7 | 14 |
| 2015 | PN | 159 | 173 | 222 |  |  |
| 2016 | PNP | 171 | 180 | 215 | 0 | 14 |
| 2016 | PN | 171 | 180 | 229 |  |  |
| 2017 | PNP | 151 | 163 | 200 | 0 | 11 |
| 2017 | PN | 151 | 163 | 211 |  |  |

### Global view of gene expression patterns

We harvested triplicate samples in 2014 and 2017 from flowering until after veraison (for time points see Figure 1 and Additional file 1: Tables S2 and S3), individual harvests are referred to below as subsamples, and analyzed them by RNA-Seq. After preprocessing the raw data (see Methods), reads derived from each subsample were mapped to the reference sequence from PN40024 and analyzed with respect to the CRIBI V2.1 annotation dataset. For 2014, approximately 19.7 million reads per subsample were obtained from each of the 72 libraries. An overall alignment rate of 79 % to the grape reference genome sequence was reached. For 2017, approximately 43.5 million reads per subsample were obtained from each of the 78 libraries. From these, an overall alignment rate of 92 % to the reference was calculated. Expression values were initially detected as Transcripts Per Kilobase Million (TPM) and averaged over the three subsamples for each sample. Considering both years separately, a total of 28,692 genes were detected as expressed in both cultivars and in both years. In contrast, 2,152 CRIBI V2.1 genes were found to be not expressed.

The correlation between gene expression data, determined as TPM values per sample, of the datasets from both years over all genes was r = 0.5095 (Pearson correlation coefficient) for PN and r = 0.6557 for PNP, respectively. For PN and PNP, 10,205 and 16,226 genes, respectively, expression values were significantly correlated (p-value < 0.05) between the years 2014 and 2017. A list of the correlation strength of the eight time points with the same BBCH stage is provided in Additional file 1: Table S4.

To visualize global trends and similarity of the gene expression values obtained from all subsamples, a Principal Component Analysis (PCA) of both years was performed with keeping the subsamples separate. The first component PC1 explains 57% of the variance, while the second component PC2 explains 17% (Figure 2). Almost all data points of the subsamples (triplicates within a sample) from both years cluster near to each other. The data follow a track of time in a nearly consecutive and continuous way. Main actors, which influence most of the variance in the data, were genes related to cell wall modification, secondary metabolism, wounding-response and hormone signaling. The top 500 genes responsible for most of the variance in PC1 and PC2 are listed in Additional file 1: Table S5 and Table S6, together with functional information for each listed gene from MapMan/Mercator and RefSeq.

**Figure 2:**
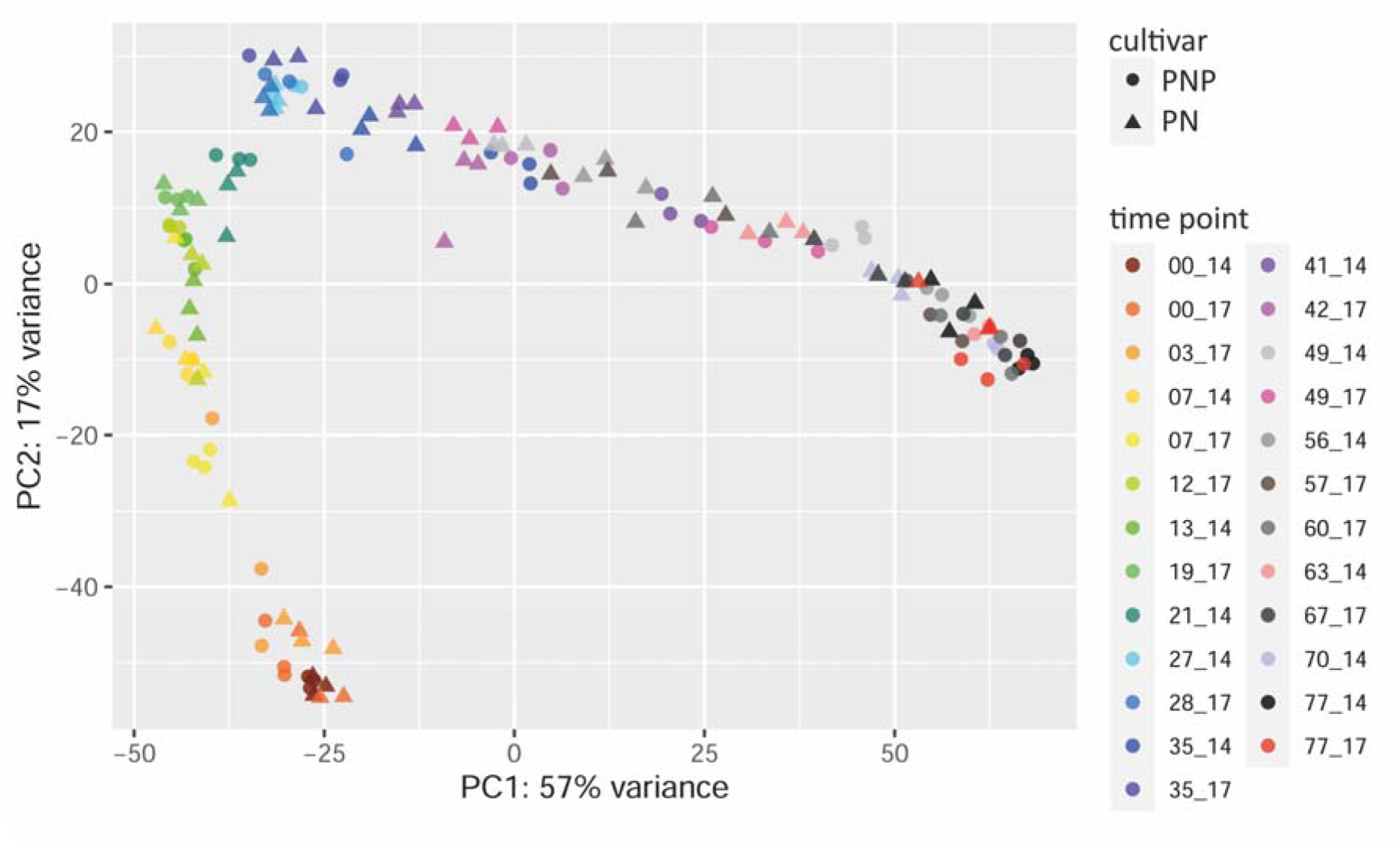
Principle Component Analysis (PCA) of gene expression values from all subsamples. Each data point represents a single subsample of the triplicates for each time point of both years (2014 and 2017 as indicated by [DAF]_14 and [DAF]_17 with the colour code) and for both cultivars (PN as triangles, PNP as circles).

### Cluster analysis for identification of co-expressed genes

Gene expression time series profiles combined data from the subsamples/triplicates for each time point across two years for both culrivars. Expression profiles were compared using the clustering tool CLUST. The goal was the characterization of similarity and/or differences in gene expression among years and cultivars throughout berry development. Over all four datasets, 13 PN/PNP clusters of genes with similar gene expression patterns (C1-C13) were obtained (Additional file 2: Figure S1A). In these clusters, 3,316 (12.2 %) of the 27,139 genes expressed during berry development were found co-expressed among both years and cultivars. It should be noted that CLUST uses criteria to define expressed and not expressed genes that differ from the ones applied above (see Methods). The observed expression profiles differ clearly between the clusters, which was in part the result of the restricted number of clusters that CLUST extracts. Manual inspection of the clusters revealed little deviation of individual gene expression profiles within each individual cluster for a given year or cultivar. All cluster gene memberships, also those for the additional cultivar-specific cluster analyses (see below), are available in Additional file 1: Table S7-S9.

The PN/PNP clusters C2, C5, C6 and C12 (C12_PN/PNP selected as example, see Figure 3) reveal a small but detectable difference in the gene expression profile between both sampled years, but are almost identical for both cultivars. The PN/PNP clusters C1, C7 and C11 (C1_PN/PNP selected as example, see Figure 3) show similar expression profiles over the two years, but stand out by shifted expression peaks that distinguish PN and PNP.

**Figure 3:**
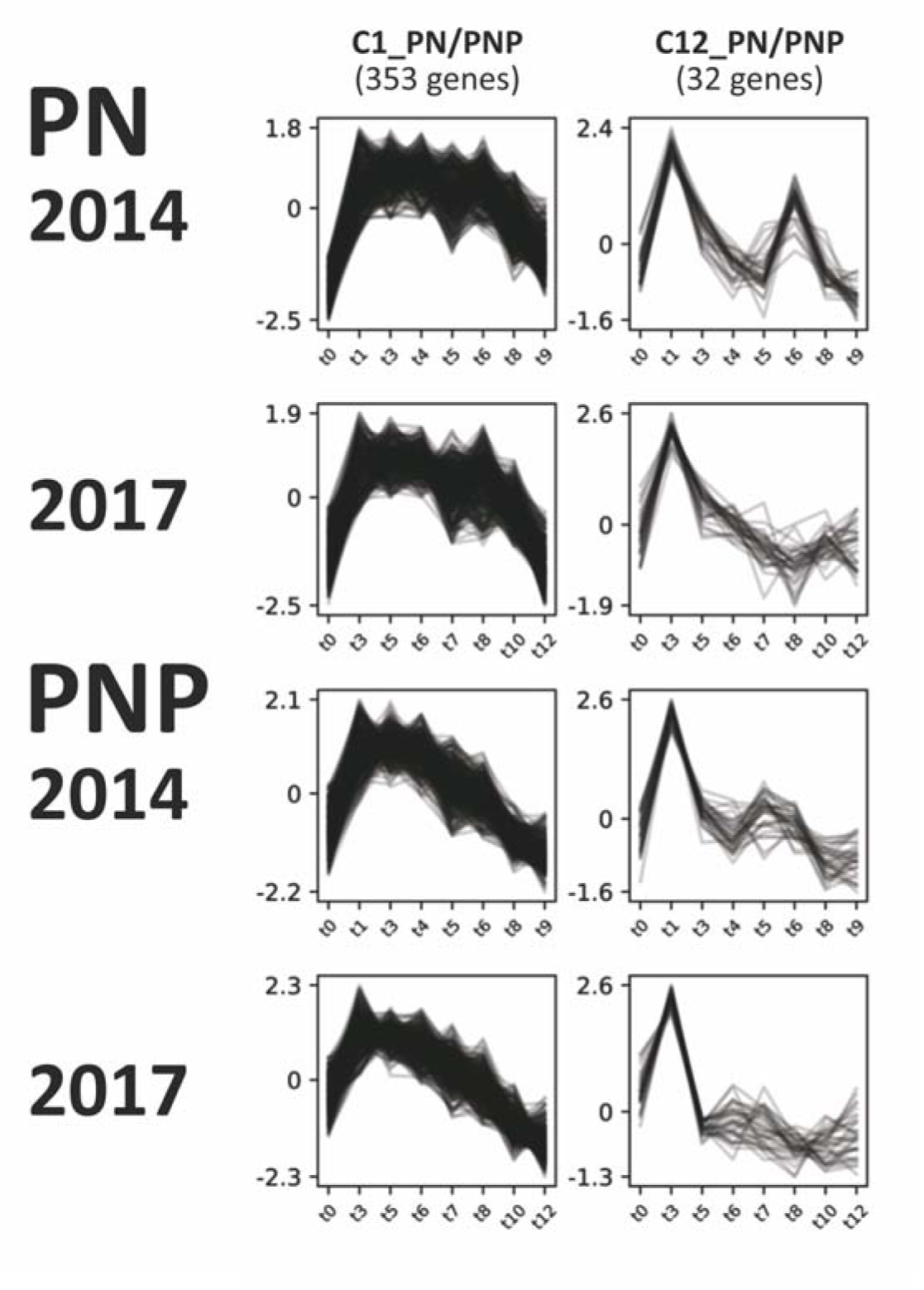
Two selected gene expression profile clusters with either a cultivar-specific difference (C1_PN/PNP) or a weather/field condition-specific difference (C12_PN/PNP) after clustering all data (both cultivars and both years). Strength of gene expression (quantile normalization) was plotted over the time course of berry development. Sampling time points are detailed in Figure 1 and were restricted to those eight equivalent time points at which the cultivars display the same BBCH stage (Additional file 1: Table S4). For all PN/PNP clusters see Additional file 2: Figure S1A. IDs of genes that make up the clusters are listed in Additional file 1: Table S7.

To characterize the clusters with respect to potential functions of the co-expressed genes included in a given cluster, GO term enrichment for biological processes was calculated. The full list of enriched GO terms for all clusters is listed in Additional file 1: Table S10-S12. Two examples for GO terms appearing with highly significant incidence were ‘response to oxidative stress’ in cluster C11_PN/PNP (term GO:0051276) and ‘regulation of defense response’ in cluster C5_PN/PNP (term GO:0031347).

Two additional cluster analyses were performed, one for the PN data from both years (Additional file 2: Figure S1B) and one for the PNP data from both years (Additional file 2: Figure S1C). These analyses revealed a high abundance of genes from the expansin gene family with similar expression profiles in clusters C0_PNP and C6_PN. The cluster C16_PN was found to have a highly significant enrichment for ‘vegetative to reproductive phase transition’ (GO:0010228). Cluster-gene memberships for the cultivar-specific clustering are available in Additional file 1: Table S8-9, and the corresponding GO term enrichment is summarized in Additional file 1: Table S11-12.

### Analyses of differentially expressed genes

The gene expression time series throughout berry development was analyzed for differentially expressed genes (DEGs) between the two cultivars PN and PNP with DESeq2. Genes with significantly differential expression were selected by using the filters adjusted p-value (PADJ) < 0.05 and log2fold change (LFC) > 2. The results are summarized in Table 2 and are detailed at the gene level per time point compared in Additional file 1: Table S13.

**Table 2:** Filtering steps applied for selecting DEGs, and the number of DEGs that were carried on after each selection step. For details see Methods.

|  | PN/PNP 2014 [DEGs] | PN/PNP 2017 [DEGs] |
| --- | --- | --- |
| **adjusted p-value (PADJ) < 0.05:**<br>(counted over all sample pairs) | 8,206 | 4,419 |
| **log2fold change (LFC) > 2:**<br>(counted over all sample pairs) | 6,629 | 4,298 |
| **down- / up-regulated**<br>(in PNP vs. PN): | 3,293 / 3,336 | 2,130 / 2,168 |
| **unique:**<br>(non-redundant within time series) | 3,342 | 2,745 |
| **intersection:**<br>(detected in both years) | 1,923 |  |
| **excluded due to intersection:** | 1,419 | 822 |
| **veraison-specific genes:**<br>(detected within BBCH79-81 of PNP) | 388 |  |
| **potentially regulatory:**<br>(detected within BBCH61-79 of PNP) | 12 |  |

In total, 8,206 and 4,419 DEGs were identified with PADJ greater than 0.05 for 2014 and 2017, respectively. Almost twice as many DEGs were initially detected in 2014 samples compared to 2017. By applying the filter for an at least 2-fold difference in expression level (LFC > 2), the number of significant DEGs decreased, mainly for the PN/PNP time series from 2014. Only few DEGs between PN and PNP were observed during flowering (BBCH61 to 69) at the beginning of both time series (see Figure 4). Within the berry formation phase (BBCH71 to 79), the number of DEGs detected increased towards veraison (BBCH81) as both genotypes increasingly varied in physiological stages. The highest number of DEGs was observed in parallel to the time-shifted veraison of PNP relative to PN. This time point was also the most phenotypically different between the two cultivars (see Figures 1 and 4). A set of veraison-specific genes was defined by selecting the DEGs from time points DAF35 and DAF41 from 2014 that were also observed to coincide with this phenotype at DAF42 and DAF49 in the 2017 gene expression data. These criteria identified 388 veraison-specific DEGs. This set of 388 genes was compared to results from similar studies and found to be in agreement (e.g. 81% [27] and 52% [28]; IDs of the 388 genes, the genes that match results from the other studies and their functional annotation, are included in Additional file 1: Table S14). During the subsequent phase of berry ripening (BBCH81 to 89, after around DAF56 in 2014 and around DAF60 in 2017), the number of DEGs detected decreased. We developed a visualization for the numbers of DEGs detected and the changes with respect to which genes are newly appearing as differentially expressed at a given time point (sample pair PN/PNP) in the time series (Figure 4). Groups of newly appearing DEGs are containing only few genes early in berry formation, while numbers increase at veraison of PNP. After veraison of PN, the number of DEGs decreased. If DEGs appearing in several time points are counted only once, 3,342 and 2,745 unique DEGs (different genes) are detected from 2014 and 2017, respectively (compare Table 2).

**Figure 4:**
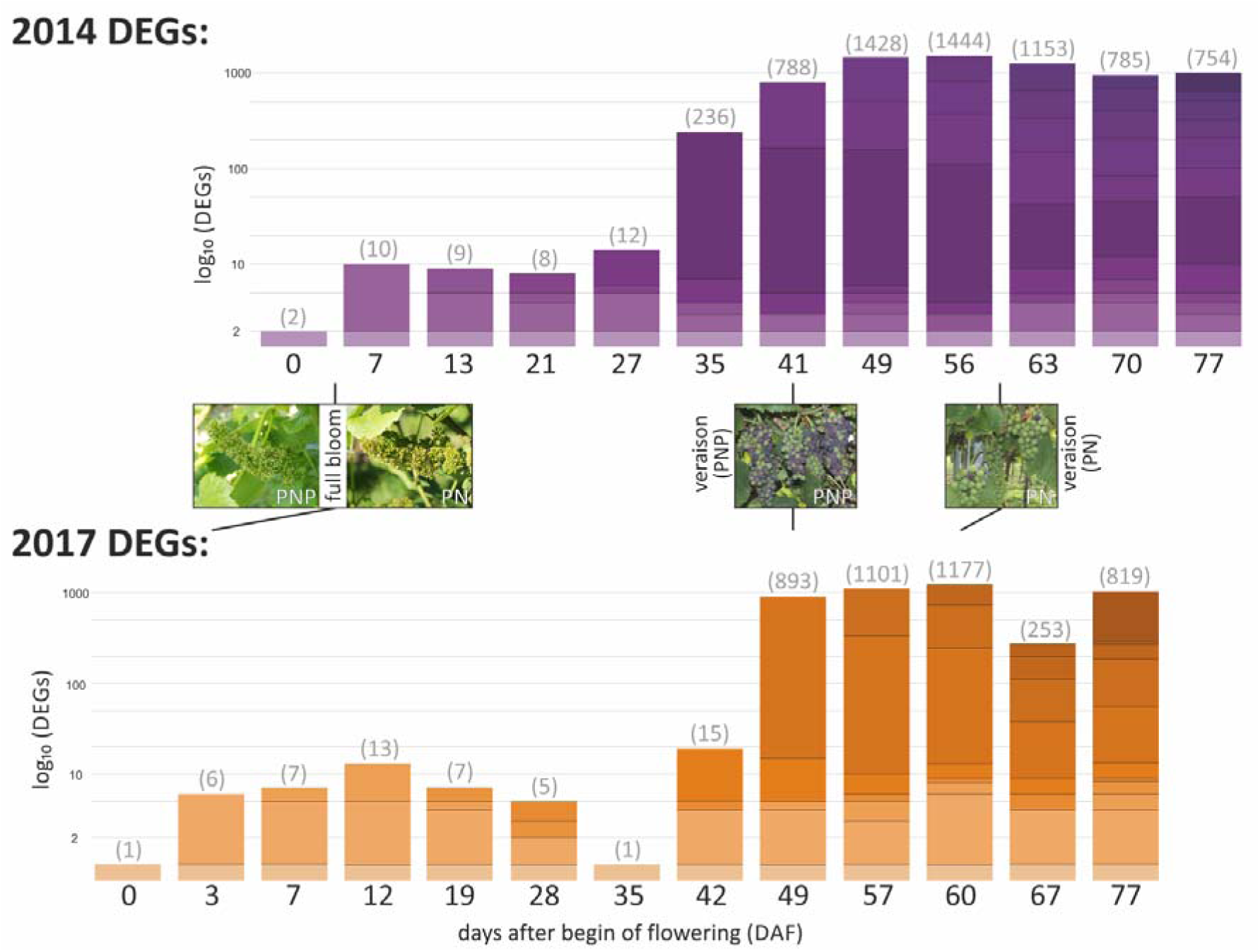
Visualization of the number of DEGs detected between PN and PNP in a logarithmic scale (log_10_). Results for 2014 are shown in purple, those for 2017 in orange. The time series from the two years were aligned at veraison of PNP; the timeline is given as days after onset of flowering (DAF). DEGs are counted for a pair of PN/PNP samples for each time point individually, the number above each column mentions the number of DEGs detected at the respective time point. Groups of newly appearing DEGs relative to an earlier time point are indicated by a new colour shade in the column (bar) for each time point. For members of a given group of DEGs, the attributed colour shade is kept for the subsequent time points (columns/samples). The pictures between the two column series display BBCH65 (full bloom, 50% of flowerhoods fallen [3]) and BBCH81 (veraison) of PNP and PN.

To further increase the reliability, reproducibility, and relevance of the selected DEGs, the intersection between the DEGs identified in the two years studied was computed. In total, 1,923 unique DEGs were obtained (Table 2). To reveal DEGs potentially involved in the control of timing of ripening, i.e. genes that might be involved in the trait that mainly distinguishes PN and PNP, only intersecting DEGs which appeared at time points before veraison in PNP were picked. This resulted in a list of 12 DEGs that may control ripening time. It should be noted that these putative regulatory DEGs are supposed to be relevant before the set of veraison-specific genes implements the phenotypic changes at veraison. The full list of DEGs, their identity and annotation information as well as their fit to the selection criteria on the way from all (raw) DEGs to potentially regulatory DEGs are detailed in Additional file 1: Table S13. IDs of the 12 putative ripening time control genes, the genes that match results from related studies (7 DEGs [27], 4 DEGs [34] and 3 DEGs [28]) and their functional annotation are included in Additional file 1: Table S14, the most relevant data are summarized in Table 3.

**Table 3:** Collection of features of the set of 12 genes classified as potential regulators based on their differential expression before veraison of PNP. The detailed expression patterns are shown in Figure 5 and Additional file 2: Figure S3. Details on annotation are listed in Additional file 1: Table S13.

| gene ID | gene symbol | functional annotation<br>(transferred via BLASTp) | relative transcriptional change |  |
| --- | --- | --- | --- | --- |
|  |  |  | pre veraison<br>of PNP | before, at or after veraison<br>of PN & PNP |
| VIT_205s0051g00590 | *VviPL1* | pectate lyase 8 | upregulated in PNP | peaks before veraison, goes down after veraison |
| VIT_205s0077g01980 | - | uncharacterized protein LOC100248252 | upregulated in PNP | peaks before veraison, stays up after veraison |
| VIT_206s0004g00070 | *VviEXPA5* | expansin A10 | upregulated in PNP | upregulated in PN, strong peak before veraison |
| VIT_206s0009g02560 | *VviPME10* | pectinesterase 2 | upregulated in PNP | off in PN, may come up later after veraison |
| VIT_209s0018g01490 | - | methanol O-anthraniloyltransferase | upregulated in PNP | goes up before veraison, further up after veraison |
| VIT_210s0071g01145 | *VviRTIC1* | plant protein with Domain of Unknown Function 789 | upregulated in PN | off in PNP |
| VIT_213s0067g02930 | *VviEXPA14* | expansin A8 | upregulated in PNP | upregulated in PN, peak before veraison, down after |
| VIT_216s0013g00880 | *VviOLE5* | oleosin 1 | upregulated in PNP and PN | peaks before veraison, low consistency between years |
| VIT_216s0022g00960 | *VviGRIP28* | ripening-related protein-like precursor | upregulated in PNP | goes up before veraison, stays up after veraison |
| VIT_216s0098g01170 | *VviHDZ28* | homeobox-leucine zipper protein ATHB-12 | upregulated in PN | up after flowering, goes down long before veraison |
| VIT_200s0366g00020 | *VviRTIC2* | cysteine-rich receptor-like protein kinase 10 | upregulated in PN | off in PNP |
| VIT_200s0956g00020 | *VviLEC1* | nuclear transcription factor Y subunit B-6 | upregulated in PNP and PN | peaks before/at verais., low consistency between years |

### Functional classification of DEGs

To complement the gene lists with functional information from grapevine that might potentially be informative for berry development, the 1,923 intersecting DEGs were analyzed with respect to enrichment of genes that have been assigned to biological pathways already established for grapevine (see Methods). For 46 of the 247 defined grapevine pathways, significant enrichment (permuted p-value <0.1) was detected. The most reliable predictions (permuted p-value <0.001) for pathways that might be relevant were photosynthesis antenna proteins (vv10196; 9 DEGs); nitrogen metabolism (vv10910, 19 DEGs); phenylpropanoid biosynthesis (vv10940, 66 DEGs), tyrosine metabolism (vv10350, 33 DEGs); transport electron carriers (vv50105, 18 DEGs); phenylalanine metabolism (vv10360, 33 DEGs); brassinosteroid biosynthesis (vv10905, 8 DEGs) and flavonoid biosynthesis (vv10941, 30 DEGs). The enrichment results are provided in (Additional file 1: Table S15). The same analysis was also carried out for the 12 putative ripening time control (Additional file 1: Table S16) and the 388 veraison-specific DEGs (Additional file 1: Table S17).

A check of the 1,923 intersecting DEGs revealed that 141 TF genes are included. Of these, 48 DEGs were clearly up- and 93 down-regulated at their first appearance in the time series. The full list of TF encoding genes that were higher expressed in PNP (up-regulated), or lower expressed in PNP (down-regulated), compared to PN, is shown in Additional file 1: Table S18. For a more detailed view on the expression patterns of selected TF encoding genes, we generated for the TF gene family with the highest abundance among the 141 TF genes, namely the R2R3-MYB-type TFs with 22 cases in the MapMan functional assignment, an expression heatmap (Additional file 2: Figure S2). As a result, *VviMYB24* (VIT_214s0066g01090), which is related to At3g27810/*AtMYB21*, At5g40350/*AtMYB24* and At3g01530/*AtMYB57* according to TAIR/PhyloGenes, was identified as an early appearing DEG that showed its highest expression level at flowering (BBCH61). Prominent *R2R3-MYB* genes known to be relevant for anthocyanin accumulation like *VviMYBA1* (VIT_202s0033g00410), *VviMYBA2* (VIT_202s0033g00390), *VviMYBA3* (VIT_202s0033g00450) and *VviMYBA8* (VIT_202s0033g00380) were detected as expressed starting from veraison (BBCH81) in both cultivars and with a time shift towards earlier expression in PNP. An additional *R2R3-MYB* gene with a similar expression pattern is *VviMYB15* (VIT_205s0049g01020). Other *R2R3-MYB* genes are expressed early during berry formation, these include *VviMYBF1* (VIT_207s0005g01210, related to At2g47460/*AtMYB12*/*AtPFG1*) as well as *VviMYBPA5* (VIT_209s0002g01400) and *VviMYBPA7* (VIT_204s0008g01800, both related to At5g35550/*AtMYB123*/*AtTT2*). According to their related expression patterns visualized in the heatmaps (Additional file 2: Figure S2), the *R2R3-MYB* genes fall into three groups that roughly fit to the three phases marked in Figure 1B, namely flowering, berry formation, and berry ripening (see discussion).

### Putative candidates for ripening time control genes

As mentioned above, DEGs detected in both years at time points before veraison of PNP were selected and considered as putative genes that control ripening time (Table 3, Additional file 1: Table S13). The VitisNet enrichment analyses performed for these 12 candidates resulted in 2 pathways that showed significant (permuted p-value < 0.05) enrichment with two genes in the pathway: auxin signaling (vv30003 with *VviEXPA5* (VIT_206s0004g00070) and *VviEXPA14* (VIT_213s0067g02930)) and cell wall (vv40006 with *VviPL1* (VIT_205s0051g00590) and *VviGRIP28* (VIT_216s0022g00960)); see Additional file 1: Table S16).

A detailed check of the data presented in Figure 4, together with results for these putatively ripening time control DEGs, resulted in the identification of two DEGs that stand out from the whole list of DEGs. Both genes almost completely lack expression in the early ripening cultivar PNP while there is clear expression in PN. Therefore, these two genes were detected as DEGs throughout the whole time series in both years. The first of the two, designated *VviRTIC1* for “Ripening Time Control” (VIT_210s0071g01145, encoding a protein similar to “protein of unknown function DUF789”), is expressed during flowering (BBCH61 - 65) and is more or less continuously down-regulated over time in PN (Figure 5A). The second of the two, designated *VviRTIC2* (VIT_200s0366g00020, encoding a protein similar to “cysteine-rich receptor-like protein kinase”), displays expression in PN during berry formation as well as during berry ripening with a peak before veraison in 2017 (Figure 5B). The expression patterns of the 10 remaining DEGs of the putative ripening time control gene set are shown in Additional file 2: Figure S3.

**Figure 5:**
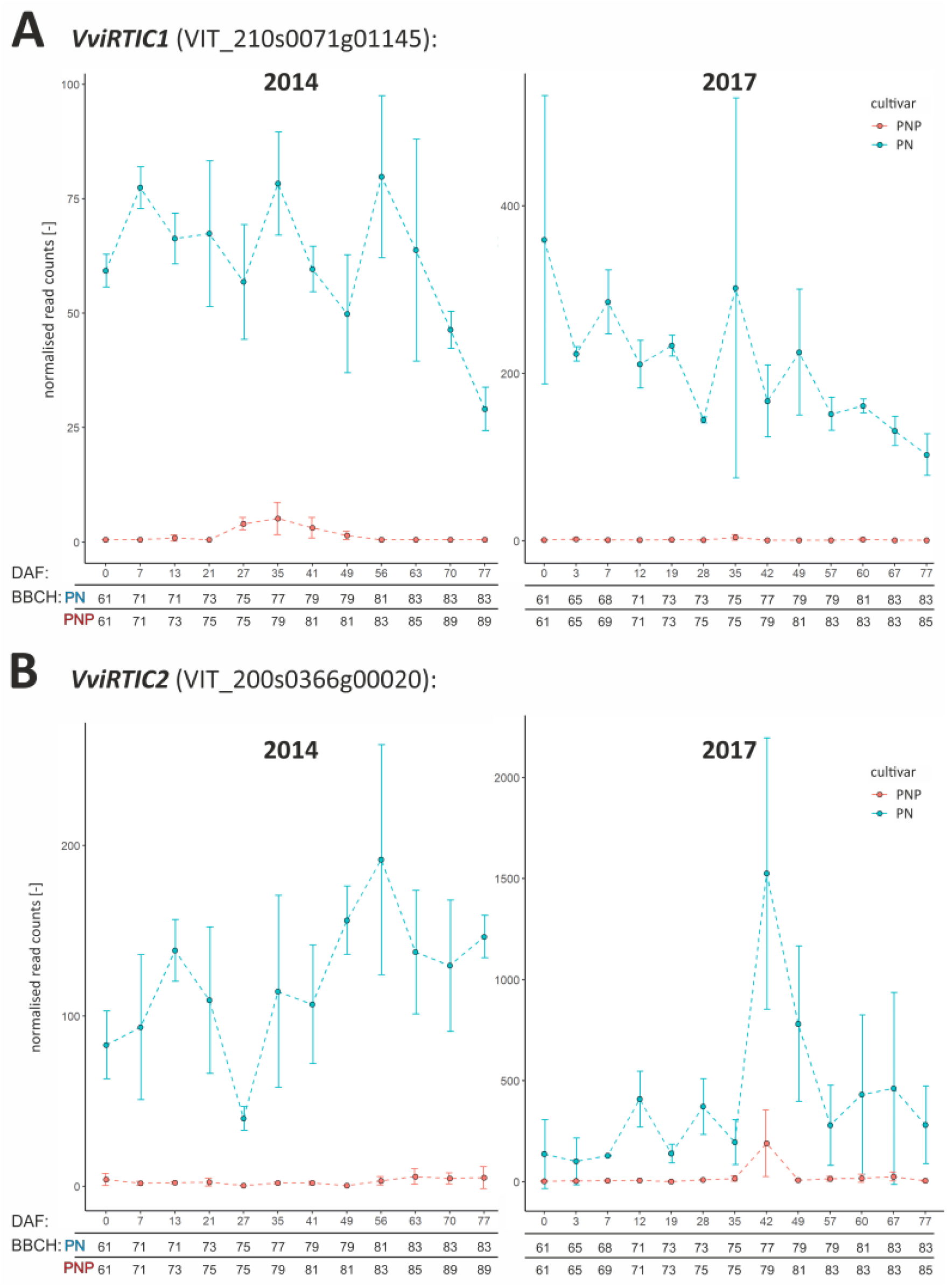
Expression patterns of *VviRTIC1* (VIT_210s0071g01145) in (A) and *VviRTIC2* (VIT_200s0366g00020) in (B) from RNA-Seq data of PN (blue) and PNP (red). Error bars display the standard deviation of triplicates. Left, expression profile from 2014. Right, expression profile from 2017. The y-axis represents the read counts from the output of DESeq2. The x-axis represents the development stages in days after onset of flowering (DAF).

Data on the set of 12 putative ripening time controlling genes are collected in Table 3, with a focus on relative up- or downregulation of expression before veraison of PNP. Also, a short description of the differences of the expression patterns before, at and after veraison of PN and PNP is included.

The expression patterns derived from RNA-Seq confirm each other and also allow digital quantification of low transcript accumulation levels. Nevertheless, three genes were chosen for confirmation via qRT-PCR, namely *VviRTIC1*, *VviRTIC2* and *VviERF027* (VIT 216s0100g00400). *VviERF027* was included to cover a gene that displays, in different samples of the time series, differential expression as well as equally high expression in PN and PNP. The results obtained by qRT-PCR are fully congruent with the data from RNA-Seq (see Additional file 2: Figure S4). Thus, differential gene activity in a developmental pattern and in a genotype specific way has been detected by RNA-Seq as well as qRT-PCR.

## Discussion

One of the first detectable mentions of the cultivar ‘Pinot Precoce’ in connection with the synonym ‘(German) Früh Burgunder Traube, translated: Early Burgundy Grape’ (PNP) is in the French book “Ampelographie retrospective” [37]. PNP is considered to be closely related to ‘Pinot noir’ (PN) grapes and, here, we have confirmed the clonal relationship of PN and PNP by 24 well distributed genomic SSR markers. Although this does not prove that PN is the ancestor, it is very likely that PNP was derived from the cultivar PN by somatic mutation as suggested earlier [36]. We used these two isogenic cultivars, that are distinguished by a clear duration of berry formation phenotype, to analyze changes in gene expression throughout berry development. The aim was to identify candidate genes that control the speed of berry development and veraison timing. Samples from inflorescences as well as from forming and ripening berries were collected from the onset of flowering until after PN and PNP veraison in 2014 and 2017. These samples were subjected to RNA-Seq analyses in two well resolved time series.

### Phenotypic differences between the cultivars PN and PNP

The data from 4 years of careful assessment of the BBCH developmental stages of PN and PNP at the same location validate earlier observations from viticulture [36] that lead to establishment of PNP as a distinct grapevine cultivar in north European wine growing countries. Berry formation lasts about two weeks less in PNP, is clearly accelerated compared to PN and results in PNP entering veraison approximately two weeks earlier than PN (Figure 1A, Table 1). It is reasonable to assume that this acceleration affects berry formation throughout, from immediately after fruit set until veraison. Functionally, this hypothesis implies that the genes that are responsible for the control of timing of berry development, and for the establishment of the phenotypic difference between PN and PNP, should be acting already very early in berry development, starting at least shortly after flowering and at or even before BBCH61 to BBCH79. At the end of ripening (harvest), the berries of PNP reach high sugar content earlier within the season when compared to PN.

### General validation of the RNA-Seq dataset

To estimate overall data quality, the expression profiles obtained from PN and PNP were correlated for the two sampled years, 2014 and 2017. Pearson correlation was moderate, but this is expected considering the conditions of the free field environment. Exposure of the vines to external factors like biotic or abiotic stressors, including weather conditions that differ significantly between the years, also affect the transcriptome which reduces the level of correlation [38]. In a PCA, almost all data points lie on the same intended track, and biological replicates (subsamples) from both years are located close to each other. The main actors, which predominantly influence the variance in the dataset, are genes related to cell wall modification, secondary metabolism, wounding and hormone signaling. These gene categories fit locical expectations since berry development is known to (i) be controlled by hormones, (ii) require new cell walls, and (iii) be accompanied by accumulation of secondary/specialized metabolites [24, 26]. These initial results validated the quality of the dataset and indicated clearly that sampling of biologically closely related material for the subsamples/triplicates was successful.

### Co-expression analysis shows similar gene expression clusters between cultivars and years

To further validate the data with respect to comparability as well as reproducibility between the two years, related gene expression profiles were identified among all genes by clustering the data from the four different time series. Generally, clusters of the same genes with similar expression patters over time were observed for both cultivars and both years. Also, the cluster analyses for gene expression patterns among the years 2014 & 2017 in only PN and in only PNP, confirmed comparability of the gene expression patterns obtained in these two years. Detailed inspection revealed clusters representing expression profiles (and clusters of genes) with and without an environmental influence. Especially the cultivar-specific clusters C1_PN/PNP, C7_PN/PNP and C11_PN/PNP stand out. Comparison of the expression profiles for PN to those of PNP in these clusters identified a similar pattern that is moved to a different time in PNP. These findings are in agreement with the shifted ripening time phenology of the two cultivars discussed above. In contrast, the clusters C2_PN/PNP, C5_PN/PNP, C6_PN/PNP and C12_PN/PNP display more pattern similarity among the two years than among the two cultivars. Thus, the genes in these four clusters may display dependence on environmental factors in their expression patterns, potentially due to differences in the weather conditions between the two years studied. Strong environmental effects on gene expression patterns have also been described for grapevine berry development at 11 different environments (vineyards) from northern Italy [39]. The remaining other PN/PNP clusters C0, C3, C4, C8, C9 and C10 display highly similar expression profiles ofer al four conditions. The genes included in these clusters are probably less affected by environmental factors and/or the genotypic difference between PN and PNP. We conclude that our RNA-Seq results and expression level comparisons between two years are based on valid data.

When the genomic location of the DEGs is analyzed, a genome region on chromosome 16 comes into focus. In this region, 54 of the DEGs from the set of 1,923 intersecting DEGs (Table 2) are located. Of these, 28 encode stilbene synthases [40] that are all up-regulated after veraison of PNP (BBCH83). Stilbenes are a group of phenylpropanoid compounds (that includes resveratrol) which are detected in many plants, which often accumulate in response to biotic and abiotic stresses, and which are formed as a basic structuce by the key enzyme stilbene synthase. The genome region fits to a major QTL (Ver1) for “timing of the onset of veraison” on chromosome 16 [8]. It remains to be determined if this aggregation of DEGs is by chance. Potentially, the observation is biased by co-regulation of a large number of closely linked stilbene synthase genes.

### Differentially expressed genes throughout berry development and identification of veraison-specific genes

Differential gene expression analysis and subsequent filtering revealed 1,923 DEGs between PNP and PN. DEG detection was based on a comparison of samples taken from the two cultivars at very similar DAF. As expected for the characterized phenotype, PNP reaches veraison when PN is still in the phase of berry formation. Consequently, the strong increase in the number of detected DEGs shortly before and at veraison of PNP results from the different developmental stage of PNP compared to the lagging PN. Subsequently, when also PN enters veraison, the number of DEGs declines (note that Figure 4 uses a logarithmic scale). A list of 388 genes that show up in both years with a veraison-specific expression pattern was extracted and compared to published results. Interestingly, about 81.5% of the 388 PN/PNP veraison-specific genes were also described in the 4,351 differentially expressed genes between the table grape cultivar ‘8612-CTR’ (wild type) and its early ripening bud mutation ‘8612-ERN’ [27]. Also, analyses of berries from the cultivars ‘Cabernet Sauvignon’ and ‘Pinot Noir’ by RNA-Seq identified a gene set of 5,404 genes marking the onset of berry ripening [28]. This set covers 51.5% of the 388 PN/PNP veraison-specific genes (Additional file 1: Table S14). Several “switch genes” which are supposed to encode key regulators of the developmental transition at veraison [34, 41] are also included in the 388 veraison-specific gene set (Additional file 1: Table S14). We conclude that the PN/PNP veraison-specific set of 388 genes represents a core set of genes that are relevant for executing the switch from berry formation to berry ripening. As a relatively small gene set was detected that still displays high overlap to those found by other studies that addressed similar biological questions, indicates that the specific experimental setup and implemented filters used here are appropriate to remove unrelated genes. Here, the comparison of “wildtype to mutant” RNA-Seq results in isogenic background between PN and PNP, reduced environmentally controlled transcriptome differences by sampling in the same vineyard/location, and dense time course sampling together with high RNA-Seq read coverage allowed good resolution power.

In order to check for potentially co-expressed genes within the veraison-specific gene set, the memberships of these genes in the PN/PNP cluster analysis were investigated. A total of 48 veraison-specific DEGs were detected in cluster C6_PN/PNP (contains 914 genes). These 48 genes include several prominent ripening-related genes like *VviGRIP61* (VIT_201s0011g05110), *VviMYBA8*, *VviMRIP1* (VIT_205s0049g00760, [42]), *VviGRIP4* (VIT_205s0049g00520) and *VviGRIP28*. The ∼20 *VviGRIP* genes were previously detected by differential cDNA screening as ripening-induced genes in grape [43]. Another relevant cluster is C5_PN/PNP (contains 263 genes) which includes 37 of the 388 veraison-specific DEGs. Among these are *VviMYBA1*, *VviMYBA2*, *VviMYB15* and *VviGRIP22* (VIT_206s0004g02560).

The two clusters C5_PN/PNP and C6_PN/PNP show quite similar patterns (Additional file 2: Figure S1A). It was, at first, not obvious which difference has forced CLUST to put a given gene in either C5_PN/PNP or C6_PN/PNP. However, a comparison of the expression patterns of *VviMYBA2* (in C5_PN/PNP) and *VviMYBA8* (in C6_PN/PNP; see Additional file 2: Figure S2 for a heatmap) shows that there are borderline cases regarding assignment to either C5_PN/PNP or C6_PN/PNP.

In total, 22 genes encoding R2R3-MYB TFs were found among the 1,923 intersecting DEGs. Based on the timing of expression in PN and PNP, the 22 *R2R3-MYB* genes can be classified into three groups (Additional file 2: Figure S2). The first group is represented by *VviMYB24* which is expressed during early flowering (BBCH61) but switched off already at the transition from flowering to berry formation (BBCH71). *VviMYB24* is potentially orthologous to a group of three *A. thaliana R2R3-MYB* genes (*AtMYB21*/*24*/*57*) that are expressed in flowers and which function redundantly to regulate stamen development in the context of jasmonate action [44]. It is tempting to speculate that *VviMYB24* has a similar function in grape.

The second group covers about 15 *R2R3-MYB* genes that are expressed during berry formation and pre-veraison (BBCH71 to 77). This group includes *VviMYBF1* which regulates flavonol biosynthesis [45], and *VviMYBPA5* as well as *VviMYBPA7* which belong to the clade of *AtTT2*-related genes that control proanthocyanidin (PA, flavan-3-ol) biosynthesis [7, 46–48]. The other *R2R3-MYB* genes in this group are less well characterized although there are functions described for some of them, e.g. *VviMYBC2-L3* (VIT_214s0006g01620) as repressor of specific branches of the phenylpropanoid pathway [49].

The third group of *R2R3-MYB* genes is active starting at veraison (after BBCH81) and covers about six genes. Among them are the anthocyanin accumulation controlling genes, *VviMYBA1*, *VviMYBA2, VviMYBA3* and *VviMYBA8* for which there is good evidence that they trigger anthocyanin biosynthesis [50]. Since PN and PNP are red berry cultivars, activity of the TF genes that direct anthocyanin accumulation is expected. In addition, this group includes *VviMYB14* and *VviMYB15* that are supposed to regulate the stilbene biosynthetic pathway [51]. With regard to the heatmaps (Additional file 2: Figure S2) and the analyses of the DEGs in this study in general, it should be noted that while the resolution within the developmental program and time is quite good, our data do not resolve the exact location of gene expression. Therefore, it remains to be determined if the expression detected is derived from berry skin, flesh, the seed or other tissues/cells.

### Putative ripening time control genes acting early in berry development

To focus on genes that are contributing to the acceleration of berry formation in PNP, and/or to the control of timing of veraison, we selected DEGs detected at time points prior to veraison of PNP (Table 2, Figure 4). This resulted in a set of only 12 genes that are potentially involved in the regulation of ripening time (Table 3). According to our hypothesis that the genes relevant for acceleration of berry formation in PNP, which cause the earlier onset of ripening in PNP, should be acting from at least shortly after flowering, we designated this set of genes as “ripening time controlling”. However, genes that encode components of the respective regulatory networks and target genes of regulators including secondarily affected DEGs are surely included as well [52]. The 12 putative ripening time control genes, the DEGs detected before veraison of PNP, encode proteins related to auxin action, pectin processing enzymes related to cell wall modification, TFs from the HD-Zip as well as NF-Y/LEC families, a cysteine-rich receptor-like protein kinase, an oleosin, and proteins with domains of unknown function.

The expression patters of this set of differentially expressed genes, and also the complete dataset of DEGs detected (Table 2), was screened for genes that were higher expressed in PNP than in PN. However, although there are many genes that display earlier upregulation of expression in PNP than in PN in the context of earlier veraison, none of the 12 “early differential” putative ripening time control genes was significantly higher expressed in PNP than in PN already at or early after flowering (Table 3). Also, among the other DEGs, no gene with such a differential expression pattern was detected. Such early gene activation in PNP compared to PN could hint at a dominant regulator that promotes faster ripening, but the data are more in favor of loss of an inhibitor of fast ripening.

The two genes assigned to auxin signaling by VitisNet (vv30003) encode expansins (*VviEXPA5* and *VviEXPA14*, [53]). Expansins are known to be involved in fruit ripening through cell wall expansion and cell enlargement [54]. Auxin can delay the onset of veraison and ripening processes in grapevine [19–21]. Since reduced expression of genes from the auxin signaling pathway may indicate reduced auxin action due to lower auxin levels, the accelerated entry of PNP into veraison might be initiated by reduced auxin levels. Additionally, the genes *VviPL1* (pectate lyase 1 [55]), *VviPME10* (pectin methylesterase 10, VIT_206s0009g02560) *VviGRIP28* (encoding a pectin methylesterase inhibitor precursor-like protein) are also related to cell wall processes, indicating that cell wall modification is an important target process also prior to veraison [54]. The gene *VviGRIP28* was also detected within a veraison-specific meta-QTL designated ver/ph16.1 [9]. It remains to be determined if this correlation has a functional basis.

The two genes in the set of 12 that encode TFs are *VviHDZ28* (VIT_216s0098g01170, [56]) and *VviLEC1* (VIT_200s0956g00020, [57]). The *V. vinifera* gene VIT_216s0098g01170 that has been designated *VviHDZ28* has also been considered as a homolog of *AtHB12* (At3g61890), but it seems that *VviHDZ07* (VIT_202s0025g02590) and *VviHDZ27* (VIT_215s0048g02870) are more similar to *AtHB12*. In these cases, which lack clearly assignable homologs, transfer of functional information reaches its limits and might be restricted to concluding that *VviHDZ28* is important for organ development in *Vitis*. The gene *VviLEC1* is one of three genes in *V. vinifera* which are homologs of *AtLEC1* (At1g21970, NF-YB9) and *AtL1L* (LEC1-like, At1g21970, NF-YB6). LEC1 and L1L are central regulators of embryo and endosperm development. They control, among other processes, embryo morphogenesis and accumulation of storage reserve [58]. It is tempting to speculate that the reason for the detection of *VviLEC1* among the 12 putative ripening time control genes is that also seed development needs to be accelerated in PNP compared to PN. This would explain earlier and higher expression of *VviLEC1* in PNP compared to PN as observed (Additional file 2: Figure S3G). Consequently, *VviOLE5* (VIT_216s0013g00880, encoding an oleosin involved in oil body formation [59]) would fit into the picture as relevant for lipid storage during seed development. According to the proposed enzyme function as an alcohol acyltransferase by the protein encoded by VIT_209s0018g01490 involved in volatile ester formation [60], this gene could play a similar role. For the gene VIT_205s0077g01980 no functional annotation is available (uncharacterized protein), although homologs exist throughout the Magnoliophyta.

### Candidates for causal genes explaining the difference between PN and PNP

Among the 12 putative ripening time control genes, of which 10 are discussed above, two are especially interesting. Detailed analyses of the full set of DEGs, visualized in Figure 4, resulted in the identification of *VviRTIC1* and *VviRTIC2*, that could possibly be centrally involved in the accelerated berry development and earlier beginning of ripening in PNP compared to PN. The special feature of the expression patters of the two genes (Figure 5, Table 3) is that both are differentially expressed already at the first time point analyzed which was selected to hit the BBCH stage 61 (flowering before full bloom, DAF zero (0)). Also, both genes are only barely expressed in PNP in both years studied, while expression in PN is high at almost all time points. *VviRTIC1* is annotated to encode a protein containing a domain of unknown function (DUF789), while *VviRTIC2* is annotated to encode a “cysteine-rich receptor-like protein kinase”. The best BLASTp hit to *A. thaliana* protein sequences indicates that it is related to At4g23180/AtCRK10, but a closer inspection shows that similarity to At4g05200/AtCRK25, At4g23160/AtCRK8 and At4g23140/AtCRK6 is almost as high. This ambiguity, and also the fact that the *V. vinifera* genome contains several genes related to *VviRTIC2* (e.g. VIT_210s0071g01200, VIT_202s0087g01020 or VIT_203s0017g01550 as listed by PhyloGenes), complicates transfer of functional information. CRKs are a subgroup of plant receptor-like kinases [61] and are encoded by a family of 44 genes in *A. thaliana*. In a systematic analysis of the functions of *A. thaliana* CRKs, evidence was collected for involvement in the control of plant development, biotic and abiotic stress responses, photosynthesis as well as stomatal regulation [62]. This systematic phenotypic screen of a large set of T-DNA insertion mutants revealed distinct phenotypes for various of the CRK genes, but assignment of a molecular function to individual CRKs beyond recognition of unknown ligands and signal transmission by phosphorylation remains a large challenge.

As pointed out above, it is very possible that the genes we have identified are part of a genetic pathway that controls timing of berry development in *V. vinifera*, but that we have hit genes in a downstream part of this pathway. The relevance of the two candidate genes in the causal genetic difference between PN and PNP remains to be determined. Phase-separated genome sequences of the cultivars will be required to resolve the genome structure of both alleles of *VviRTIC1 and 2* the genes in PN and PNP for an informative comparison. In future studies, we will address this question, for example by long read DNA sequencing.

## Conclusions

This study detected 1,923 DEGs between the Pinot cultivars PN and PNP. The two clonal cultivars display a phenotypic difference in berry development timing where PNP reaches from full bloom to veraison faster than PN. We defined 388 DEGs as veraison-specific and 12 DEGs as putatively controlling ripening time. The relatively small number of veraison-specific genes displays a very high overlap with results published for similar studies (see Additional file 1: Table S14) and could be used for studying a phytohormone network that is acting similarly in PN and PNP, but accelerated in PNP. Additionally, the ripening time control genes identified here might offer access to a set of genes putatively important for triggering or delaying the start of berry ripening in grapevine. Further investigations are needed to elucidate structural differences in the genomes, the function of the observed DEGs, and their role in shifting the onset of ripening in grapevine.

## Material and Methods

### Plant material and analysis of clonal relation

The grapevine (*Vitis vinifera* subsp. *vinifera* L.) cultivar PNP (Pinot Precoce Noir, VIVC No. 9280) is early ripening and has been described to be related to the cultivar PN (Pinot Noir, VIVC No. 9279) [35] that ripens later than PNP. To prove the clonal relation, DNA from both cultivars was genotyped utilizing 24 polymorphic SSR markers (VVS2, VVMD7, VVMD5, VVMD32, VVMD28, VVMD27, VVMD25, VVMD24, VVMD21, VVIV67, VVIV37, VVIQ52, VVIP60, VVIP31, VVIN73, VVIN16, VVIH54, VVIB01, VrZAG83, VrZAG79, VRZAG67, VrZAG62, VMC4F3.1, VMC1B11) as described [63]. The two cultivars used have been identified as accession DEU098_VIVC9280_Pinot_Precoce_Noir_DEU098-2008-076 and DEU098_VIVC9279_Pinot_Noir_DEU098-2008-075, respectively. The tissue used for harvest is indicated below and in Figure 1. Both cultivars do not belong to an endangered species and were obtained and are grown in accordance with German legislation.

### Phenotypical characterization and sampling of plant material

Plant material was harvested from PN and PNP grapevines trained in trellis. The plants are growing at the vineyards of JKI Geilweilerhof located at Siebeldingen, Germany (N 49°21.747, E 8°04.678). The grapevine plants were planted with an interrow distance of 2.0 m and spacing of 1.0 m in north-south direction. Inflorescences, developing and ripening berry samples of PNP and PN for RNA extraction were collected in two years with three independent biological replicates (subsamples) each. Sampling took place at systematic time points (12 time points in 2014, 13 time points in 2017), and at approx. 8 a.m. each day. In 2014, harvesting took place regularly every 7 days with only two exceptions (one day deviation, DAF 13 and DAF 27). In 2017, harvesting was adapted to BBCH stages (Figure 1A). The timeline in both years is described as days after onset of flowering (DAF), with onset of flowering defined as the day at which 10% of the individual flowers have lost their caps (BBCH61 [3]). For each subsample within the triplicates, material from two neighboring grapevines was selected. Grapevine plants were weekly phenotyped according to BBCH stage [3, 4]. Phenotyping was performed repeatedly to ensure sampling from vines of the same development stage (e.g. percentage of open flowers during flowering, or berry development stage) to reach uniform subsamples. The phenotypical observations were summarized in Additional file 1: Table S2 and S3. From these, the durations of flowering, berry formation and berry ripening as well as the resulting shifts between the cultivars were calculated (Table 1). Furthermore, images from berry developmental stages of both cultivars were taken in 2014 for 35, 41, 49 and 56 DAF. The sampled material was directly frozen in liquid nitrogen and stored at −70°C until RNA extraction.

### RNA extraction and cDNA library construction

Biological replicates, i.e. the subsamples, were ground separately under liquid nitrogen. Total RNA was extracted using an RNA Isolation Kit (Sigma-Aldrich Spectrum™ Plant Total RNA) according to suppliers’ instructions. Quality control, determination of RIN numbers [64] and estimation of the concentrations of all RNA samples was done on a Bioanalyzer 2100 (Agilent) using RNA Nano 6000 Chips. For RNA-Seq, 500 ng total RNA per subsample were used to prepare sequencing libraries according to the Illumina TruSeq RNA Sample Preparation v2 Guide. For subsamples from 2014 and 2017, 72 and 78 libraries were constructed and sequenced, respectively. Enrichment of poly-A containing mRNA was performed twice, using poly-T oligos attached to magnetic beads included in the Illumina kit. During the second elution of the poly-A+ RNA, the RNA was fragmented and primed for cDNA synthesis. After cDNA synthesis, the fragments were end-repaired and A-tailing was performed. Multiple indexing adapters were ligated to the ends of the cDNA fragments and the adapter ligated fragments were enriched by 10 cycles of PCR. After quality check using Bioanalyzer 2100 HS-Chips (Agilent) and exact quantification by Quant-iT PicoGreen dsDNA assay on a FLUOstar Optima Plate Reader (BMG LABTECH), the libraries were pooled equimolarly.

### RNA-Seq

Single end (SE) sequencing of the pooled barcoded libraries from 2014 was performed on an Illumina HiSeq1500 in High Output mode generating 100 nt reads. For samples from 2017, sequencing was done using an Illumina NextSeq500 generating 83 nt SE reads; two runs were performed with the same pool of barcoded libraries from 2017.

### Processing of RNA-Seq read data

Raw reads were trimmed with Trimmomatic (version 0.36) [65]. For raw reads from the year 2014, the following settings were used: LEADING:10 TRAILING:10 SLIDINGWINDOW:4:15 MINLEN:50. In addition, a collection of all available Illumina adapter sequences was supplied to remove matches within the parameter 2:30:10. For raw reads from the year 2017, trimming settings were set to LEADING:6 TRAILING:6 SLIDING WINDOW:4:15 MINLEN:36. All trimmed reads were quality-checked via FastQC (version 0.11.8) [66]. Thus, possible adapter sequences and low-quality bases were removed. All trimmed reads passing QC were mapped to the reference genome sequence PN40024 (version 12Xv2) [67] using the graph-based alignment tool HISAT2 (version 2.1.0) [68, 69] with no additional soft clipping. Afterwards, all tagged genes (structural gene annotation: CRIBI v2.1) were counted as raw read counts with FeatureCounts (Bioconductor package Rsubread version 3.8 [70]). To estimate transcript abundance as a measure for gene expression, counts for Transcripts Per Kilobase Million (TPM, [71]) were determined.

### Basic gene expression analyses

TPM counts from the various samples were used for manual gene expression inspection, for determination of the number of expressed and not expressed genes, and to calculate the correlation between gene expression values from both years. Genes with a TPM value > 0 added up over all samples from one year were classified as expressed, conversely genes with a TPM value = 0 added up over all samples as not expressed. A custom python script was applied utilizing the function pearsonr from SciPy python package (v. 1.2.3) [72], which calculates the Pearson correlation coefficient and the p-value for all year-to-year comparisons. Expression data pairs for TPM counts per gene from both sampled years, averaged over the three subsamples of each sample, were used. To test correlations and relationships between expression values from the two years, where samples were harvested with slightly different sampling patterns (see Figure 1A), eight equivalent time points with the same BBCH stages between the cultivars of each year were selected (see Additional file 1: Table S4).

### Principal component analysis

To explore data similarity, a Principal Component Analysis (PCA) was calculated over all gene expression values from both years and cultivars for all subsamples. All data points were normalized using variance stabilizing transformation function ‘vst’ from the R package DESeq2 (v. 1.12.4) [73]. Subsequently, the principal components were generated using ‘prcomp’ from the R package ‘stats’ (v. 3.5.2) [74]. The resulting PCA object, displaying the main components PC1 and PC2, was plotted and exported. Additionally, genes with the highest variance contribution to PC1 and PC2 were extracted separately.

### Functional annotation of genes

Transfer of annotation information from other plant species, mainly *A. thaliana,* was calculated using MapMan’s sequence annotation tool Mercator (v. 3.6) [75, 76]. Additionally, all open reading frame (CDS from *V. vinifera*/grapevine genes) sequences were aligned to the non-redundant protein sequence data base RefSeq [77] with the basic local alignment tool for proteins BLASTp [78] (e-value ≤ 0.001). Short descriptions of gene functions were extracted and added to the gene lists in Additional file 1: Table S5, S6, S13, S14.

GO term enrichment for biological processes was calculated via the R package ‘topGO’ (v. 2.38.1) [79]. Subsequently, statistical reliability was calculated using Fishers exact test. All Gene IDs and their corresponding GO terms were extracted from the CRIBI database (http://genomes.cribi.unipd.it/DATA/V2/annotation/bl2go.annot_with_GO_description.txt). All results of the GO term enrichment are deposited in Additional file 1: Table S10-12.

### Cluster analysis

To reveal co-expressed genes over all four datasets, the tool CLUST (v. 1.10.8) was used with default parameters [80]. As input, raw read counts from eight time points were used. These time points were selected to cover the same BBCH stages of PN and PNP from the years 2014 and 2017 (Additional file 1: Table S4). First, all data were pre-processed as described in the CLUST manual. Values from corresponding subsamples (triplicates) were combined and averaged. To filter out uninformative (very low) gene expression values, an additional filter was applied: genes not reaching a sample expression value > 1 in at least three conditions and in at least one cultivar from one year were discarded (-fil-v 1 -fil-c 3 -fil-d 1). Afterwards, the data were quantile normalized according to the RNA-Seq defaults of CLUST. Genes showing a flat expression profile were filtered out by applying the default settings [80].

### Differential gene expression analyses

For analyses of differentially expressed genes, DESeq2 (v. 1.12.4; R Bioconductor) was employed. To test if gene expression differs significantly between two samples, the likelihood ratio test nbinomLRT, included in the DESeq2 package, was used. As input, raw read counts from all time points were used. Normalization factors and dispersion estimates were used as described [73]. The output table contained all differentially expressed genes (DEGs) and the corresponding values for baseMean, log2FoldChange (LFC), lfcSE (LFC standard error), stat (difference in deviation between the reduced model and the full model), p-value and PADJ (adjusted p-value). To focus on significantly differentially expressed genes from the DESeq2 analyses, cut-off filters PADJ ≤ 0.05 and LFC > 2 were applied.

### Confirmation of differential gene expression by qRT-PCR

To verify the RNA-Seq results, four time points from 2017 were selected (DAF 0, DAF 28, DAF57 and DAF77) for qRT-PCR. Synthesis of cDNA from the RNA subsamples was carried out with First Strand cDNA Synthesis Kit (ProtoScript® II; NEB) according to the manufacturer’s instructions. The qRT-PCR assay was performed using Luna Universal qPCR Master Mix (NEB) with a total volume of 20 µl. Sequences of the primers used are listed in Additional file 1: Table S23. Reaction products/amplicons were detected based on SYBR green via a CFX96 Real Time PCR Detection System (Bio-Rad). For each time point, three biological and three technical replicates (i.e., each subsample in triplicate) were measured. Cycling conditions after initial denaturation 2 min at 95°C: denaturation 5 sec at 95°C, annealing/extension 30 sec at 60°C, cycled 35 times. For QC, each reaction was controlled by product melt analysis (65°C - 95°C). As negative controls, no template control (NTC) and no reverse transcriptase control (-RT) were measured as well using three technical replicates. The polyubiquitin gene *VviUbiquitin1* (VIT_216s0098g01190) was used for normalization [46]. Measurements were analyzed using the CFX Maestro V.4.1.2433.1219 (Bio-Rad) by normalization via the relative quantitative ΔΔCt method.

### Selection of gene sets potentially relevant for ripening time and comparison with literature data

In order to identify gene sets from the DEGs relevant for control and implementation of ripening, an intersection between the DEGs detected at all time points between both years was built. To determine a subset of putatively ripening time control genes, the intersection between both years covering the development stages BBCH61 (onset of flowering) to BBCH79 (one developmental BBCH stage before veraison) was used (time points 2014: DAF 0-35; 2017: DAF 0-42). Furthermore, a set of veraison specific genes was defined from the DEGs detected at the intersection of development stages BBCH79 (one developmental BBCH stage before veraison) to BBCH81 (onset of ripening / veraison; time points 2014: DAF35-41; 2017: DAF42-49). To test for biological relevance of the subsets, all DEGs were screened to their occurrence in similar relevant studies [9, 27, 28, 30, 34, 41, 81, 82].

### Visualization of gene numbers newly appearing as differentially expressed

To visualize appearance of DEGs over time, a stacked bar plot script was set up using the R package ‘plotly’ (v. 4.9.2.1) [83]. Each bar represents the amount of DEGs of a given time point or condition. In order to track groups of DEGs newly appearing at a given time point throughout the following time points, the colour shade representing the group of DEGs remains the same.

### Pathway enrichment analysis

To search for possible targets in known pathways of grapevine, a pathway enrichment analysis using the tool VitisPathways [84] was performed. To achieve a reliable enrichment, 1000 permutations, a Fisher’s exact test of p < 0.05 and permuted p-value < 0.1 were set. Thus, all significant enriched pathway genes and their relations can be displayed in VitisNet [85], a specific molecular network for grapevine.

### Heatmaps

As an extension to assignment of genes to biosynthesis pathways, the genes were also filtered for annotation as coding for transcription factors (TFs). This filter was based on the annotation information transferred from Mercator and RefSeq (see above). To look at the entire family of *R2R3-MYB* TF genes, the list of *MYB* genes identified via MapMan was extended by additional grapevine *R2R3-MYB* gene family members that have been characterized [6, 7]. The *R2R3-MYB* genes detected among the intersecting DEGs were displayed in heatmaps addressing the four individual time series (2 cultivars, 2 years) using the R package ‘pheatmap’ (v. 2.1.3) [86]. Predictions for phylogenetic relationships were deduced from PhyloGenes v. 2.2 [87].

## Abbreviations

ABA: abscisic acid
DAF: days after onset of flowering
DEGs: differentially expressed genes
PNP: Pinot Noir Precoce
PN: Pinot Noir
LFC: log2fold change
PADJ: adjusted p-value
PCA: principle component analysis
TF: transcription factor
TPM: transcripts per kilobase million

## Declarations

### Ethics approval and consent to participate

Not applicable

### Consent for publication

Not applicable

### Availability of data and material

The FASTQ files containing all RNA-Seq reads (PN2014, PNP2014, PN2017 and PNP2017) have been deposited at the European Nucleotide Archive (ENA) according to the INTEGRAPE guidelines under the accession numbers PRJEB39262, PRJEB39261, PRJEB39264 and PRJEB39263, respectively (see Additional file 1: Tables S19 to S22). All scripts developed for this study are available on GitHub [https://github.com/bpucker; https://github.com/jenthein].

### Competing interests

The authors declare that they have no competing interests

### Funding

This project was funded by BMBF/PtJ (FKZ 031A3496).

### Authors’ contributions

Sampling and phenotyping were done at the JKI (Julius Kühn-Institute, Institute for Grapevine Breeding Geilweilerhof, Siebeldingen, Germany). Sequencing and data analysis were performed at Bielefeld University, Faculty of Biology & Center for Biotechnology (CeBiTec). Organization, supervision at Bielefeld University and parts of sampling were done by DH. KH, AK and FS coordinated weekly sampling, phenotyping, and image capture for documentation. Logistic work was done by KH. The design of the experiments was set up by FS, DH and KH. RNA isolation and cDNA synthesis were carried out by DH and PV. PV accomplished library preparation and sequencing. Quantitative RT-PCR was performed by JT and DH. RT and KH supervised the work at JKI Geilweilerhof. LH performed the SSR analysis. BW supervised the work at Bielefeld University. FS, KH, RT and BW acquired project funding and wrote the project proposal. All bioinformatic data analyses, creation of figures, tables and writing of the manuscript were performed by JT with the help of DH. JT and BW drafted the manuscript. All authors have read and approved the final manuscript.

## Acknowledgements

We would like to thank Boas Pucker for providing several python scripts for handling large datasets as well as helpful comments on the manuscript, Katharina Frey-Sielemann for help visualizing the DEG distributions and Wiebke Halpape for providing the GO annotation enquiry. We also thank Kyle J. Lauersen for higly valuable language editing, and Willy Keller for the excellent technical support. We are very grateful to the Bioinformatics Resource Facility support team of the CeBiTec for providing computing infrastructure and excellent technical support. This publication is based in part upon contributions from COST Action CA 17111 INTEGRAPE, supported by COST (European Cooperation in Science and Technology); the authors thank BMBF/PtJ for funding in the context of IPAS (project acronym novisys). The authors from UniBi also wish to thank the other members of the Chair of Genetics and Genomics of Plants for their support. We would also like to thank the Open Access Publication Funds of Bielefeld University for funding the publication charge.

## Supplementary material

**Additional file 1: Table S1**: Clonal relation of PN and PNP confirmed by 24 SSR markers

**Additional file 1: Table S2** - Sampling and phenotypical observations (BBCH) for 2014

**Additional file 1: Table S3** - Sampling and phenotypical observations (BBCH) for 2017

**Additional file 1: Table S4** - Pearsonr correlations of gene expression values over time between both years

**Additional file 1: Table S5** - Top 500 genes influencing principal component PC1

**Additional file 1: Table S6** - Top 500 genes influencing principal component PC2

**Additional file 1: Table S7** - Cluster memberships of clusters obtained from the clusteranalysis PN/PNP

**Additional file 1: Table S8** - Cluster memberships of clusters obtained from the cluster analysis PN

**Additional file 1: Table S9** - Cluster memberships of clusters obtained from the cluster analysis PNP

**Additional file 1: Table S10** - GO term enrichment for the clusters from the cluster analysis of PN/PNP

**Additional file 1: Table S11** - GO term enrichment for the clusters from the cluster analysis of PN

**Additional file 1: Table S12** - GO term enrichment for the clusters from the cluster analysis of PNP

**Additional file 1: Table S13** - Differentially expressed genes between PN and PNP in both years and filtering steps

**Additional file 1: Table S14** - Overlap of 12 putatively ripening time control DEGs (time points: DAF 0-35 from 2014 intersecting with DAF 0-42 from 2017) and 388 veraison-specific DEGs (time points: DAF 35+41 from 2014 intersecting with DAF 42+49 from 2017) with previous transcriptomic grapevine studies.

**Additional file 1: Table S15** - VitisNet enrichment for intersecting DEGs

**Additional file 1: Table S16** - VitisNet enrichment for regulatory DEGs

**Additional file 1: Table S17** - VitisNet enrichment for veraison specific DEGs

**Additional file 1: Table S18** - Up- & Down-regulated TFs among the DEGs

**Additional file 1: Table S19** - ENA sample identifier and metadata linked to the study for PNP RNA-Seq reads from 2014 (PRJEB39262)

**Additional file 1: Table S20** - ENA sample identifier and metadata linked to the study for PN RNA-Seq reads from 2014 (PRJEB39261)

**Additional file 1: Table S21** - ENA sample identifier and metadata linked to the study for PNP RNA-Seq reads from 2017 (PRJEB39264)

**Additional file 1: Table S22** - ENA sample identifier and metadata linked to the study for PN RNA-Seq reads from 2017 (PRJEB39263)

**Additional file 1: Table S23** - Sequences of primers used for qRT-PCR

**Additional file 2: Figure S1** - Cluster analyses of PN/PNP in both years (A), PN in both years (B) and PNP in both years (C)

**Additional file 2: Figure S2** - Heatmap of all MYB-TFs gene expression among the DEGs for PN and PNP in both years

**Additional file 2: Figure S3** - Expression patterns of genes from the ripening-regulatory gene set

**Additional file 2: Figure S4** - Confirmation of expression data from RNA-Seq by qRT-PCR

## Notes

### Competing Interest Statement

The authors have declared no competing interest.

### Summary of Updates

To clarify, changed ripening regulative genes to ripening time controlling genes. Added a new overview table of the genes (as potential regulators; table 3). Added new qRT PCR results (fig. S4). Optimised figures (fig.2, fig.4, fig.5 and fig.S3). Revision of the whole manuscript due to small corrections and language editing.

