## Additional-File2_figures-S1-S4 for "Transcriptomic analysis of temporal shifts in berry development between two grapevine cultivars of the Pinot family reveals potential genes controlling ripening time"

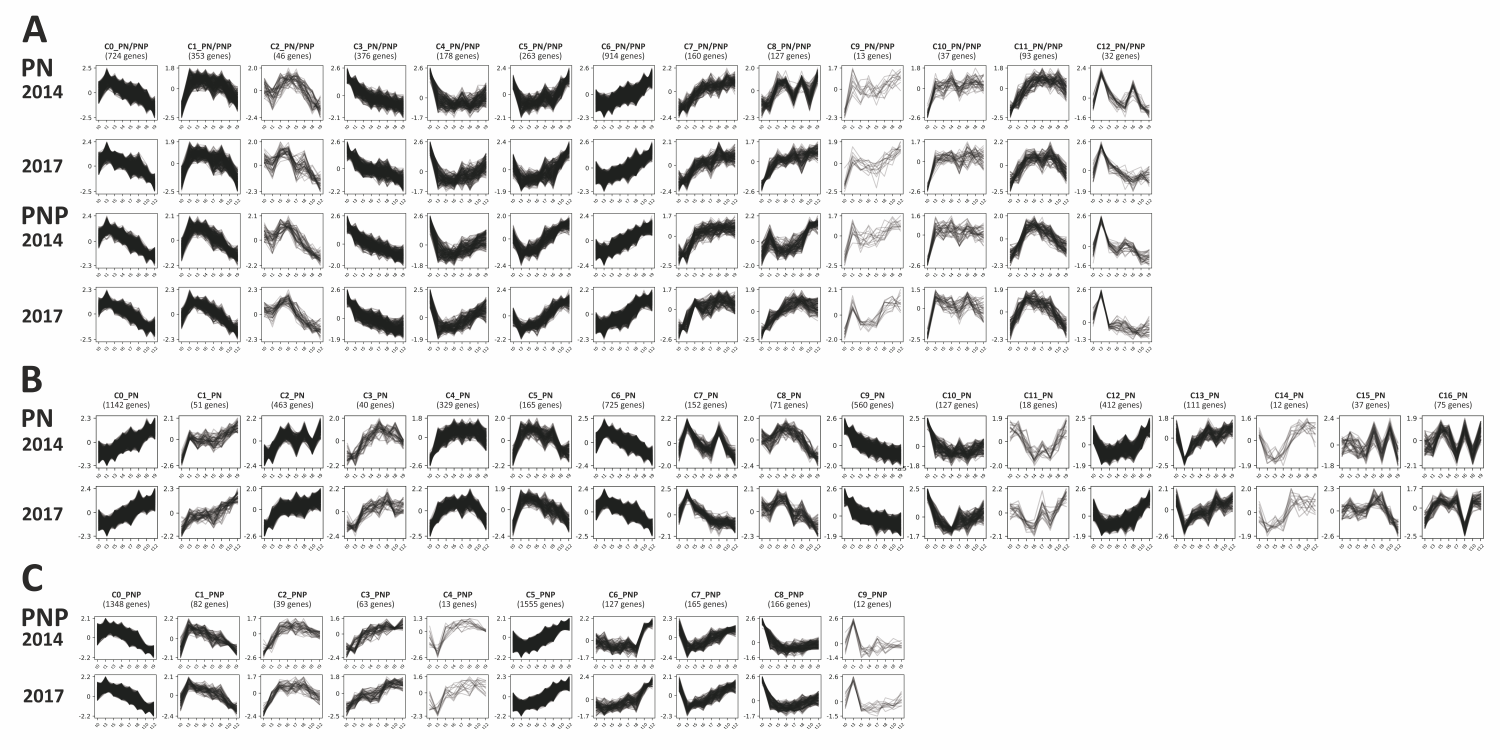

**Figure S1:** Cluster analyses of gene expression profiles. Gene expression (normalised read counts) was plotted over the time course of berry development and ripening. (A) PNP and PN in both years; (B) PNP in both years and (C) PN in both years. Sampling time points are detailed in Figure 1. IDs of genes that make up the clusters are listed in Additional file 1: Table S7 (PN/PNP), Table S8 (PN) and Table S9 (PNP).

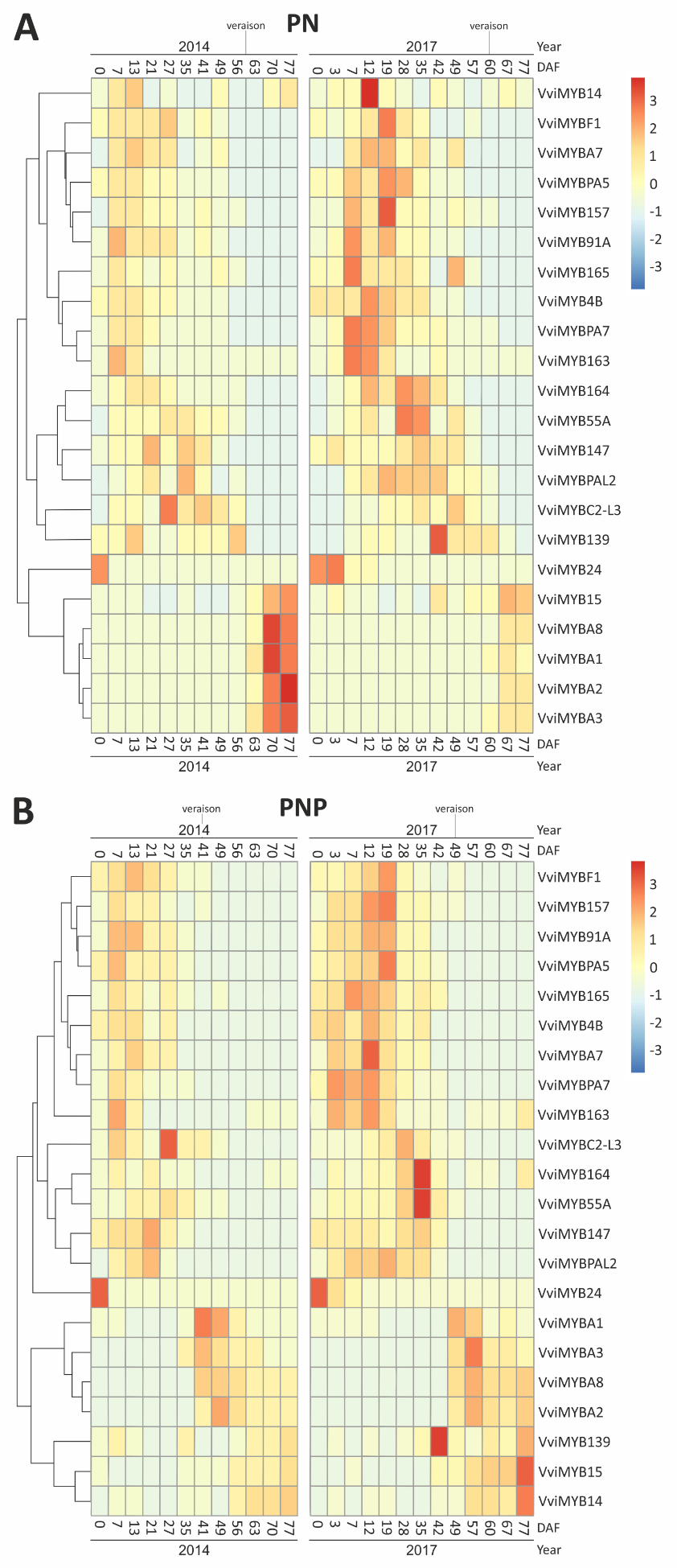

**Figure S2:** Heatmap of DEGs belonging to the R2R3-MYB transcription factor family. Expression abundance across all sampled time points of PN (A) and PNP (B) is shown for both years (2014 on the left; 2017 on the right). Relative expression levels for each gene are color-coded as indicated on the left.

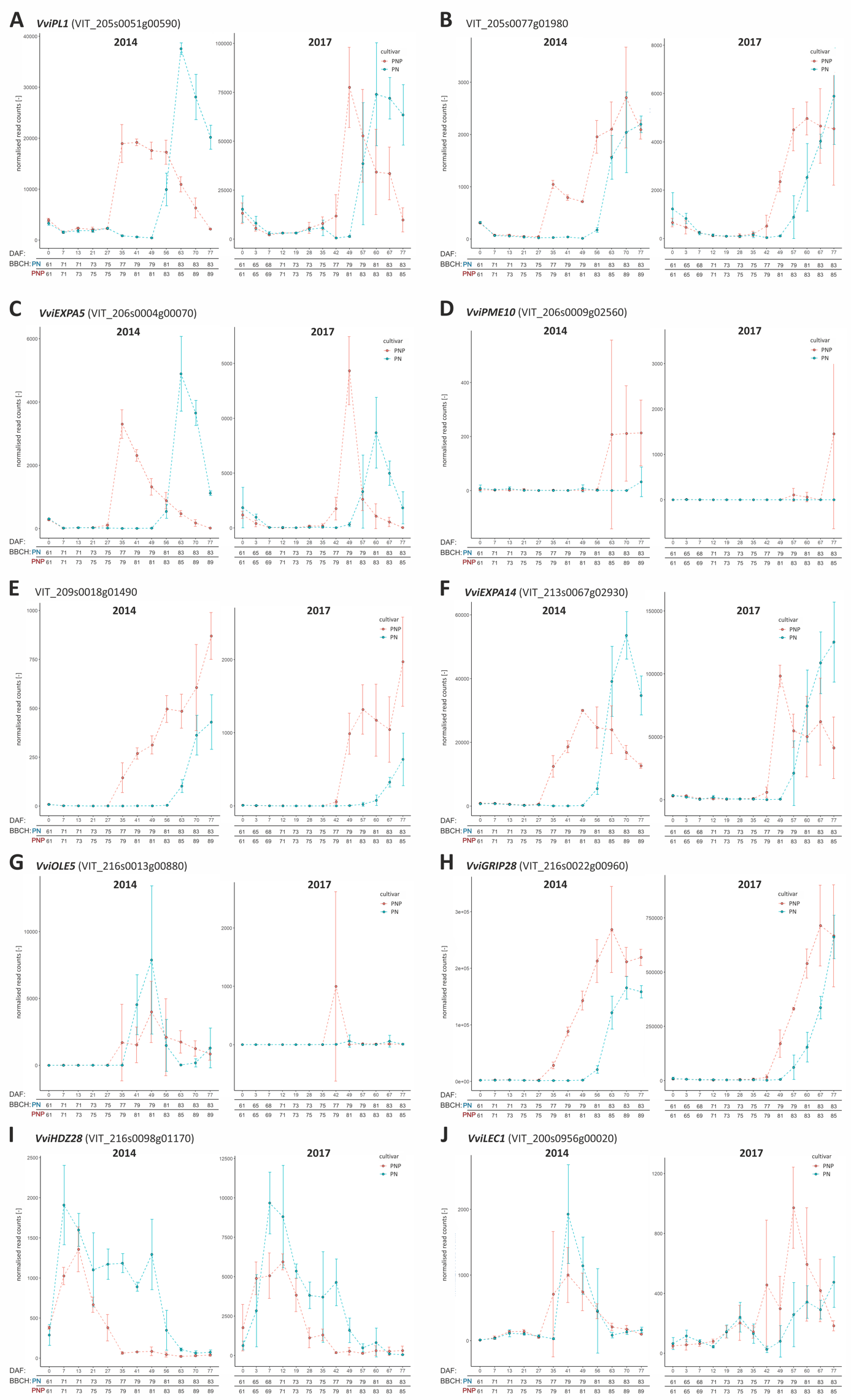

**Figure S3:** Expression patterns of the genes from the ripening-regulatory gene set, except the two shown in Figure 5.
**A**: *VviPL1* (VIT_205s0051g00590)	**B**: VIT_205s0077g01980,
**C**: *VviEXPA5* (VIT_206s0004g00070)	**D**: *VviPME10* (VIT_206s0009g02560)
**E**: VIT_209s0018g01490	**F**: *VviEXPA14* (VIT_213s0067g02930),
**G**: *VviOLE5* (VIT_216s0013g00880)	**H**: *VviGRIP28* (VIT_216s0022g00960)
**I**: *VviHDZ28* (VIT_216s0098g01170),	**J**: *VviLEC1* (VIT_200s0956g00020)
RNA-Seq data of PN are displayed in blue, and of PNP in red. Error bars display the standard deviation of triplicates. Left, expression profile from 2014. Right, expression profile from 2017. The y-axis represents the read counts from the output of DESeq2. The x-axis represents the development stages in days after begin of flowering (DAF). For inspection of *VviRTIC1* (VIT_210s0071g01145) and *VviRTIC2* (VIT_200s0366g00020) see Figure 5.

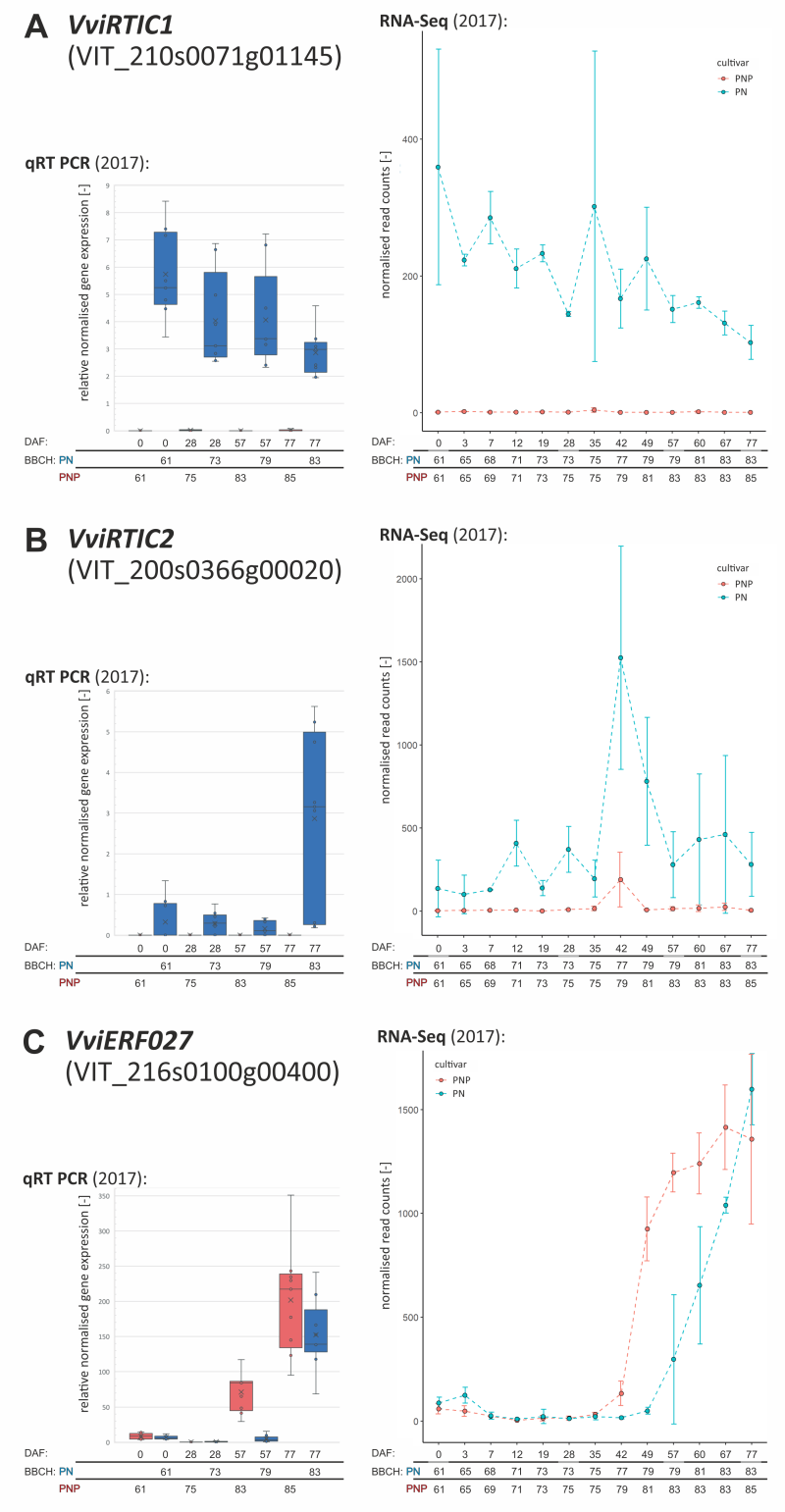

**Figure S4:** Confirmation of expression data from RNA-Seq by qRT-PCR.
Expression patterns from samples of 2017 determined by qRT-PCR (left side) and by RNA-Seq (right side). **A**: *VviRTIC1*, **B**: *VviRTIC2*, **C**: *VviERF027*. *VviERF027* was selected as veraison-specific gene from the RNA-Seq that displayed differential expression at DAF57 (no expression in PN, high expression in PNP) but equally high expression at DAF77 in PN and PNP. Profiles and data points are shown in blue (PNP) and red (PN). The x axis represents the development stages in days after begin of flowering (DAF) and their corresponding BBCH stage for each cultivar. The y axis represents the relative normalised expression from qRT-PCR and normalised read counts from RNA-Seq.
